# Comparative study of bacterial SPOR domains identifies functionally important differences in glycan binding affinity

**DOI:** 10.1101/2022.07.01.498525

**Authors:** Atsushi Yahashiri, Gabriela M. Kaus, David L. Popham, Jon C.D. Houtman, David S. Weiss

## Abstract

Bacterial SPOR domains target proteins to the divisome by binding septal peptidoglycan (PG) at sites where cell wall amidases have removed stem peptides. These PG structures are referred to as denuded glycans. Although all characterized SPOR domains bind denuded glycans, whether there are differences in affinity is not known. Here we use isothermal titration calorimetry (ITC) to determine the relative PG glycan binding affinity (*K*d) of four *Escherichia coli* SPOR domains and one *Cytophaga hutchinsonii* SPOR domain. We found that the *K*d values ranged from approximately 1 µM for *E. coli* DamX^SPOR^ and *C. hutchinsonii* CHU2221^SPOR^ to about 10 µM for *E. coli* FtsN^SPOR^. To ask whether these differences in PG binding affinity are important for SPOR domain protein function, we constructed and characterized a set of DamX and FtsN “swap” proteins. As expected, all SPOR domain swap proteins localized to the division site, and in the case of FtsN all of the heterologous SPOR domains supported cell division. But for DamX only the high-affinity SPOR domain from CHU2221 supported normal function in cell division. In summary, different SPOR domains bind denuded PG glycans with different affinity, which appears to be very important for the function of some SPOR domain proteins (e.g., DamX) but not others (e.g., FtsN).

**Importance:** SPOR domain proteins are prominent components of the cell division apparatus in a wide variety of bacteria. The primary function of SPOR domains is to target proteins to the division site, which they accomplish by binding to septal peptidoglycan. But whether SPOR domains have any functions beyond septal targeting is unknown. Here we show that SPOR domains vary in their PG binding affinities and, at least in the case of the *E. coli* cell division protein DamX, having a high-affinity SPOR domain contributes to proper function.

## Introduction

SPOR domains are typically ∼75 amino acids in length and are found in thousands of predicted proteins in various sequence databases (1). Most characterized SPOR domain proteins are involved in cell division, although there are exceptions (2–6). *Escherichia* coli has four SPOR domain proteins—DamX, DedD, FtsN, and RlpA—all of which accumulate sharply at the constriction site during cytokinesis (7–9). Septal localization of these proteins is mediated primarily by their C-terminal SPOR domains (8–10).

FtsN is the most well-studied of the four *E. coli* SPOR domain proteins. FtsN is an essential bitopic membrane protein (**Fig. 1A**) that promotes constriction by activating the septal peptidoglycan synthase FtsWI (8, 11–13). Cells depleted of FtsN form long, aseptate filaments and die. Although deletion of FtsN’s SPOR domain (FtsN^ΔSPOR^) causes a dramatic reduction in septal localization, there is only a modest cell division defect, indicating the small amount of septally-localized FtsN^ΔSPOR^ protein is sufficient to support near-normal cell division (8, 14).

**Figure 1.**
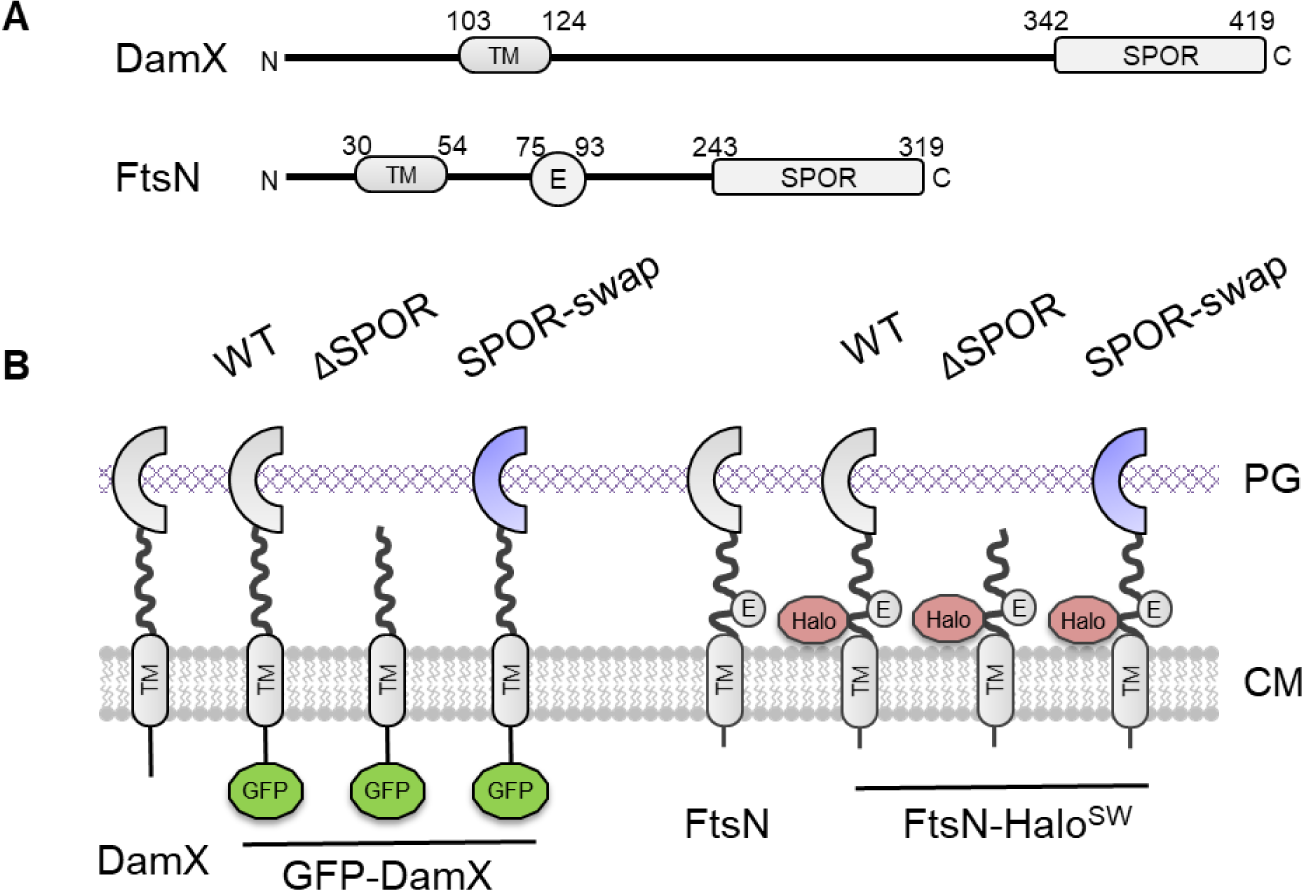
Design of DamX and FtsN SPOR domain swap constructs. (A) Domain structure of DamX and FtsN. Both are bitopic membrane proteins with an N-terminal cytoplasmic domain, a single transmembrane helix (TM), an extended periplasmic linker that is mostly unstructured, and a C-terminal SPOR domain. FtsN has in addition a periplasmic essential domain (E) that activates the FtsWI peptidoglycan synthase. (B) Swap constructs were derived from a fusion of GFP to the N-terminus of DamX (GFP-DamX) or a sandwich fusion constructed by inserting Halo between amino acids 61-62 of FtsN (FtsN-Halo^SW^).

Thus, whereas FtsN is essential for division and viability, its SPOR domain is not.

The remaining three *E. coli* SPOR domain proteins are not essential and not as well characterized as FtsN. DedD, like FtsN, is a bitopic membrane protein that promotes constriction by activating FtsWI (15). DedD null mutants have a modest elongation and cell chaining phenotype (8, 9), but DedD becomes essential when FtsN function is impaired (8) DamX is yet another bitopic membrane protein (**Fig. 1A**). DamX null mutants have no overt division defect, but they are sensitive to deoxycholate, indicative of an outer membrane (OM) integrity defect. Moreover, combining Δ*damX* with *ftsQ*(Ts) or Δ*dedD* leads to synthetic phenotypes that imply DamX has both positive and negative (inhibitory) effects on cell division (8, 9, 16). Importantly for the experiments described below, DamX’s SPOR domain is required for all of these functions (17).

RlpA is an OM lipoprotein (18). *E. coli* Δ*rlpA* mutants divide normally but deletion of *rlpA* partially rescues an *ftsK*(Ts) mutant (8, 9, 19). In *P. aeruginosa* and *Vibrio cholera*, RlpA is a lytic transglycosylase that facilitates daughter cell separation (20, 21), especially in media of low ionic strength. Curiously, *E. coli* RlpA does not hydrolyze PG and has a degenerate active site, suggesting the *E. coli* protein has lost enzymatic activity and fulfills some other function during constriction (20, 22).

Bacterial PG comprises chains of repeating β-1,4-linked *N*-acetylglucosamine (NAG) and *N*-acetylmuramic acid (NAM) disaccharides that are crosslinked by short peptides attached to the C3 of NAM residues (23). The end result is an enormous, covalently closed, bag-like macromolecule that surrounds the cytoplasmic membrane and protects against lysis due to internal turgor pressure (24). During cell division, bacteria synthesize a new cell wall through the middle of the mother cell and then carefully cleave the newly synthesized septal PG to enable daughter cell separation (25, 26). Two classes of hydrolases are particularly important for remodelling septal PG, the cell wall amidases and the lytic transglycosylases (20, 21, 27–34).

Cell wall amidases remove stem peptides from NAM residues (35). This leaves behind denuded glycan strands that serve as SPOR domain binding sites (8, 14, 36, 37). Lytic transglycosylases cleave the glycosidic NAG-NAM bonds in the amino sugar backbone of PG (35, 38) and thus destroy SPOR domain binding sites (35, 36). Importantly, the cell wall amidases and lytic transglycosylases appear to work in an ordered, sequential fashion at the septum (1, 36). The cell wall amidases act first and are the primary drivers of daughter cell separation (27, 29), while the lytic transglycosylases follow behind to clear the constriction zone of soluble PG degradation products (39).

Although SPOR domains share <20% amino acid identity in pairwise-comparisons, they adopt a conserved fold in which a 4-stranded anti-parallel β-sheet is flanked on one side by two alpha helices (17, 37, 40, 41). Mutagenesis of the SPOR domains from DamX and FtsN mapped their glycan binding sites to the β-sheet (17, 42). More recently, a crystal structure of the SPOR domain from *Pseudomonas aeruginosa* RlpA (*Pa*RlpA^SPOR^) bound to a synthetic PG glycan revealed many details of the binding interaction and explained the requirement for a denuded glycan (37). The minimal binding site is a NAG-NAM-NAG-NAM tetrasaccharide. Removal of the stem peptide from NAM leaves behind a lactyl carboxylate that makes important binding interactions with the SPOR domain β-sheet. Moreover, if the peptide were still present it would prevent binding by protruding into the SPOR domain. Interestingly, many of the SPOR-glycan contacts are mediated by main chain atoms of the SPOR domain rather than amino acid side chains. This explains how SPOR domains with very different amino acid sequences can have the same ligand specificity. Although these details were determined from the *Pa*RlpA^SPOR^-glycan complex, modeling indicates they are general features of the SPOR-glycan interaction (37).

This brings us to the questions motivating our study: Are SPOR domains interchangeable septal targeting domains? Or has evolution tailored their function to meet somewhat different needs in the context of different SPOR domain proteins? These contrasting possibilities make different predictions about what will happen when the SPOR domain from any particular protein is replaced with the SPOR domain from some other protein. Moreover, the degree of SPOR domain interchangeability might depend not only on the SPOR domain(s) but also the protein(s) to which they are attached, as these proteins might differ in how well they tolerate perturbations in the SPOR-denuded glycan interaction. We are aware of two prior reports that speak to our questions. Mӧll and Thanbichler found that a chimeric *Caulobacter* FtsN protein containing the SPOR domain from *E. coli* FtsN supported *Caulobacter* cell division (10). Taking a different approach, Gerding et al. studied the localization of GFP-tagged SPOR domains in *E. coli* and observed that partial inhibition of septal PG synthesis impaired septal localization of some SPOR domains (FtsN^SPOR^, RlpA^SPOR^) more than others (DamX^SPOR^, DedD^SPOR^, and *B. subtilis* CwlC^SPOR^) (8). These investigators suggested their findings could be explained if different SPOR domains bind different targets or if they bind to the same target but with different affinities, such that high affinity SPOR domains retain the ability to localize even when very little septal target is available.

Here we determined the relative PG glycan binding affinity for five sequence-diverse SPOR domains, which were found to be in the low micromolar range (*K*d ∼1 to ∼10 µM). We then constructed a set of DamX and FtsN SPOR domain “swap” proteins (**Fig. 1**), all of which localized sharply to division sites, as expected. Charcterization of the swap constructs in *E. coli* cell division revealed FtsN, which has the lowest affinity SPOR domain (*K*d ∼10 µM), functions well even when equipped with higher affinity SPOR domains. In contrast, DamX’s function was compromised when it’s high affinity (*K*d ∼1 µM) SPOR domain was replaced with lower affinity SPOR domains. Collectively, these results argue SPOR domains are to a large extent functionally interchangeable, but at least in the case of *E. coli* DamX a SPOR domain that binds tightly to denuded PG glycans is needed for full function.

## RESULTS

### The DamX SPOR domain binds denuded glycans in *E. coli* sacculi with a dissociation constant (*K*d) of ∼ 2 µM

We used isothermal titration calorimetry (ITC) to determine the PG binding affinity of DamX^SPOR^ from *E. coli*. ITC was suitable for our purposes because it can be used to study binding of a soluble titrant (the SPOR domain) to an insoluble ligand (PG sacculi). Moreover, ITC can be used to measure the relative *K*d knowing only the concentration of the SPOR domain but not the concentration of PG glycan binding sites, provided the latter are held constant. Our first attempts at measuring *K*d involved injecting purified DamX^SPOR^ into a suspension of purified *E. coli* PG sacculi, which were isolated from a mutant strain (EC3708) that lacks five of *E. coli’s* eight lytic transglycosylases to enrich for denuded glycans (36). Although these PG sacculi are insoluble, they were maintained in suspension during ITC by constant stirring. Injection of DamX^SPOR^ protein into the sacculi suspension produced heat changes indicative of binding, but these plateaued after just a few injections, so there were not enough data points to estimate a *K*d (**Fig. 2**). To increase the availability of SPOR domain binding sites, we subjected sacculi to partial digestion with purified AmiD amidase (43). The AmiD-treated sacculi remained largely intact and were still insoluble.

**Figure 2.**
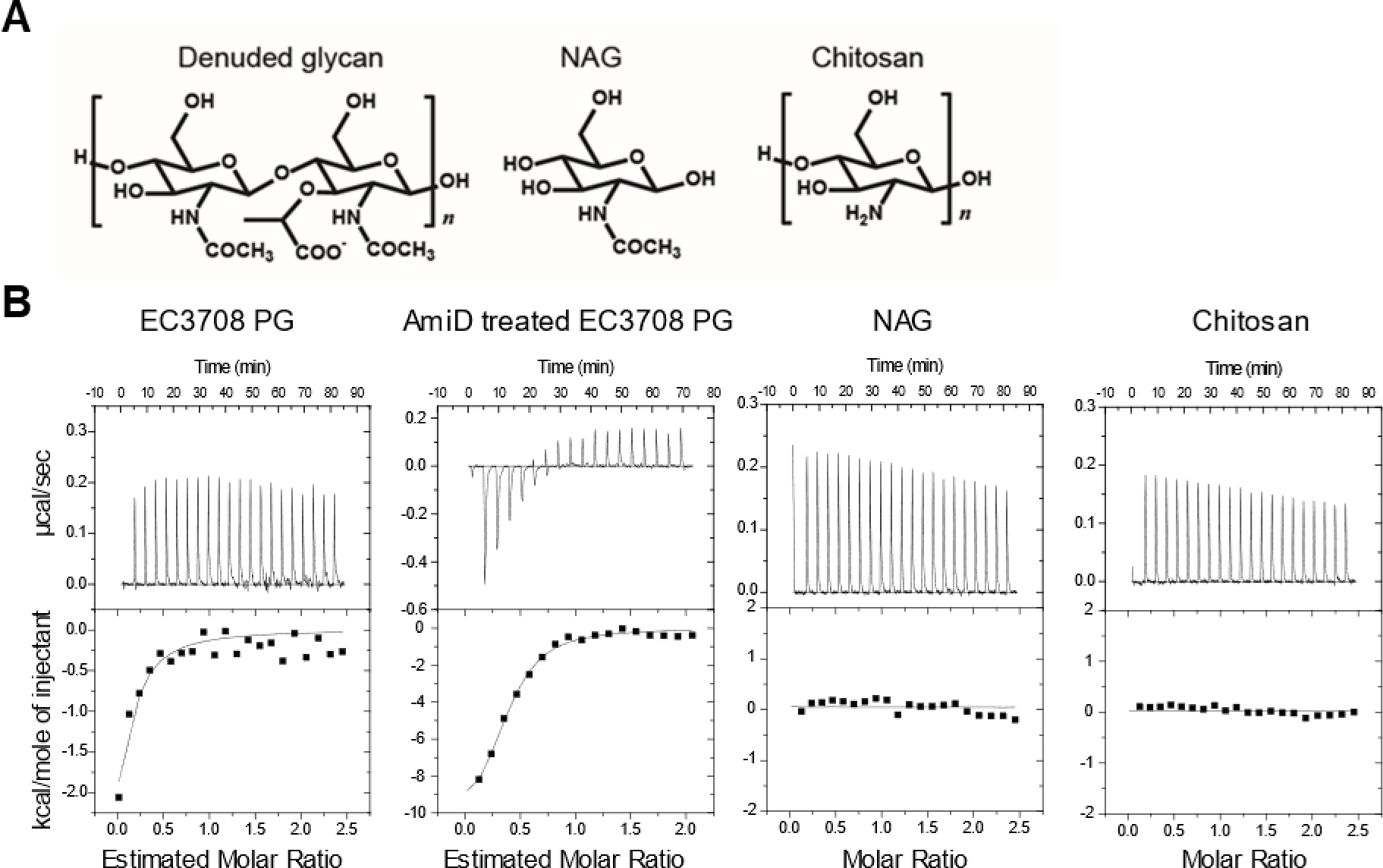
Characterization of DamX SPOR domain binding to *E. coli* PG sacculi by ITC. (A) Structure of denuded PG glycan and two analogs. (B) Representative ITC titrations showing measured heat changes as a function of time and the corresponding normalized measured heats of injection (squares) with best-fit lines. NAG, *N*-acetylglucosamine. PG was isolated from strain EC3708 (Δ*mltACDE* Δ*slt* Δ*rlpA*).

Sequential injections of DamX^SPOR^ into a suspension of AmiD-treated sacculi resulted in saturable binding over a wide-enough range of protein concentrations to determine a *K*d of 1.7 ± 0.6 µM (**Fig. 2**). No binding was observed in control experiments in which DamX^SPOR^ was injected into monomeric NAG or a chitosan polymer (**Fig. 2**).

### Multiple SPOR domains bind soluble, denuded PG glycans with *K*d’s ranging from ∼1 to ∼10 µM

We next purified and tested His6 fusions to four additional SPOR domains, three from the *E. coli* cell division proteins DedD, FtsN and RlpA, and one from an uncharacterized *Cytophaga hutchinsonii* protein named by its locus number CHU2221. All of these SPOR domains localize sharply to the division site when expressed as GFP fusions in *E. coli* (8, 9).

These SPOR domains share <20% amino acid identity in pairwise alignments (9), so we thought this set would be sufficient to uncover any differences in PG glycan binding affinity.

For denuded glycan ligand, we switched from using AmiD-treated *E. coli* sacculi to partially purified denuded PG glycans from *S. aureus*. This switch was driven by the desire to use soluble PG fragments for ITC and the practical consideration that sacculi preparations from Gram-positive bacteria yield about 10-fold more PG per liter than do those from Gram-negative organisms. *E. coli* SPOR domains localize to septal regions of PG in sacculi from *Bacillus subtilis* and *S. aureus* (1, 36), so we did not anticipate switching to Gram-positive PG would introduce any artifacts. We opted for *S. aureus* because its PG glycans are shorter and thus more soluble in aqueous buffers than *B. subtilis* glycans (23, 44, 45). After testing amidases from several organisms, we settled on the amidase domain from the major autolysin (Atl) of *S. aureus* (46, 47). The digestion products were size-fractionated by ultrafiltration to obtain soluble glycan fragments of 3-15 kDa. The glycan preparations were washed with water to remove salts, lyophilized and stored at -20°C. The final product appeared as a white, needle-like powder and could be obtained in milligram quantities from a liter of *S. aureus* culture. Based on muropeptide analysis, Atl amidase digestion removed over 90% of the stem peptides.

Nevertheless, our glycan preparations retain some peptides and are of various chain lengths. ITC titrations revealed that DamX^SPOR^ bound soluble glycans from *S. aureus* with an estimated *K*d of 1.3 ± 0.5 µM, essentially the same as obtained using AmiD-treated *E. coli* sacculi (**Fig. 2**, **Table 1**). A similar estimated *K*d of 0.8 ± 0.3 µM was obtained for CHU2221^SPOR^, whereas the SPOR domains from three additional *E. coli* cell division proteins bound less tightly: DedD^SPOR^ 3.7 ± 1.5 µM, RlpA^SPOR^ 5.1 ± 0.9 µM, and FtsN^SPOR^ 10.2 ± 2.3 µM (**Fig. S1**, **Table 1**).

**Table 1.**
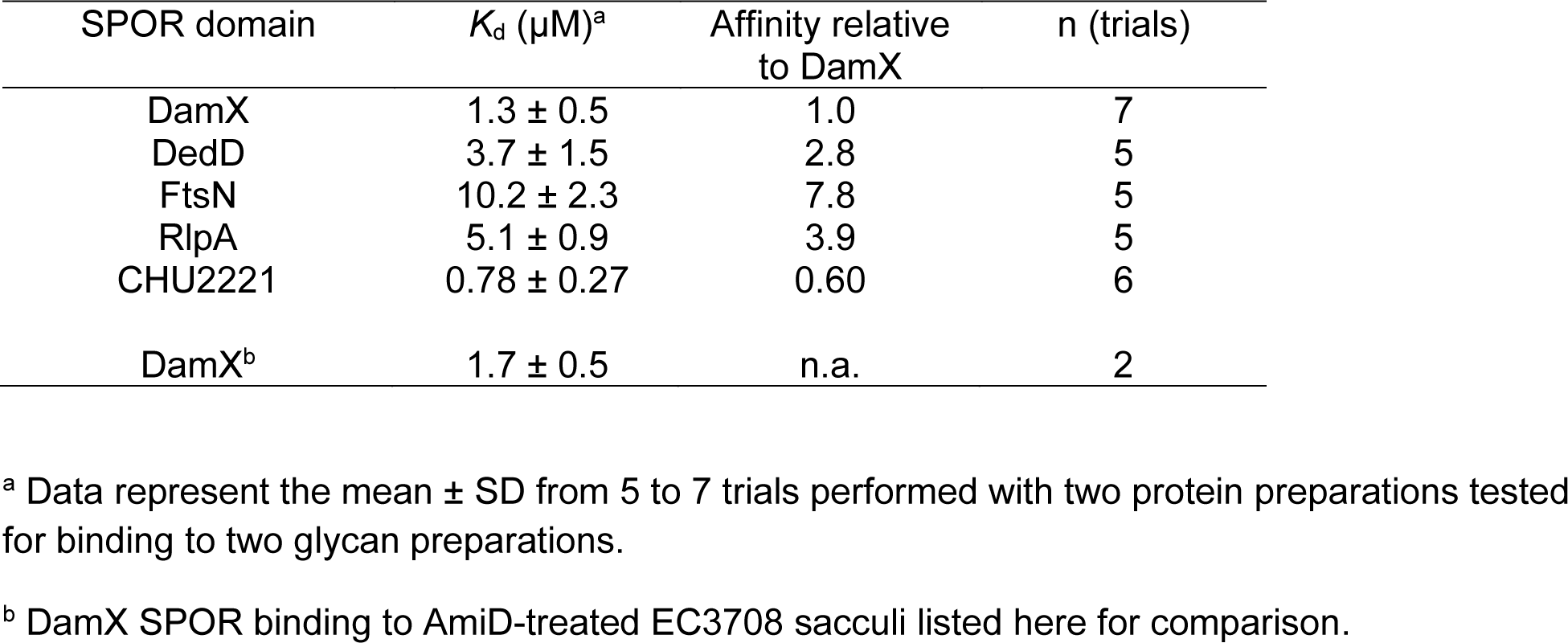
ITC characterization of SPOR domain binding to denuded glycans from S.aureus peptidoglycan

| SPOR domain | $K_d$ (μM)<sup>a</sup> | Affinity relative to DamX | n (trials) |
| --- | --- | --- | --- |
| DamX | 1.3 ± 0.5 | 1.0 | 7 |
| DedD | 3.7 ± 1.5 | 2.8 | 5 |
| FtsN | 10.2 ± 2.3 | 7.8 | 5 |
| RlpA | 5.1 ± 0.9 | 3.9 | 5 |
| CHU2221 | 0.78 ± 0.27 | 0.60 | 6 |
| DamX<sup>b</sup> | 1.7 ± 0.5 | n.a. | 2 |

<sup>a</sup> Data represent the mean ± SD from 5 to 7 trials performed with two protein preparations tested for binding to two glycan preparations.
<sup>b</sup> DamX SPOR binding to AmiD-treated EC3708 sacculi listed here for comparison.

As a control, we verified that no heat changes were produced when SPOR domains were injected into buffer (shown for DamX^SPOR^ in **Fig. S1**).

### DamX^SPOR^ and FtsN^SPOR^ compete for binding sites but DamX^SPOR^ has higher affinity

Molecular modeling predicts that DamX^SPOR^ and FtsN^SPOR^ utilize the same glycan binding sites (37). We tested this prediction using a competition assay in which mixtures of GFP-FtsN^SPOR^ and non-fluorescent DamX^SPOR^ protein were incubated with sacculi purified from an *E. coli* lytic transglycosylase mutant that accumulates denuded glycans (EC3708). After allowing for the SPOR domain proteins to bind, the sacculi were washed and visualized by fluorescence microscopy. Quantification of septal fluorescence revealed that DamX^SPOR^ reduced binding of GFP-FtsN^SPOR^ in a dose-dependent fashion (**Fig. 3A**).

**Figure 3.**
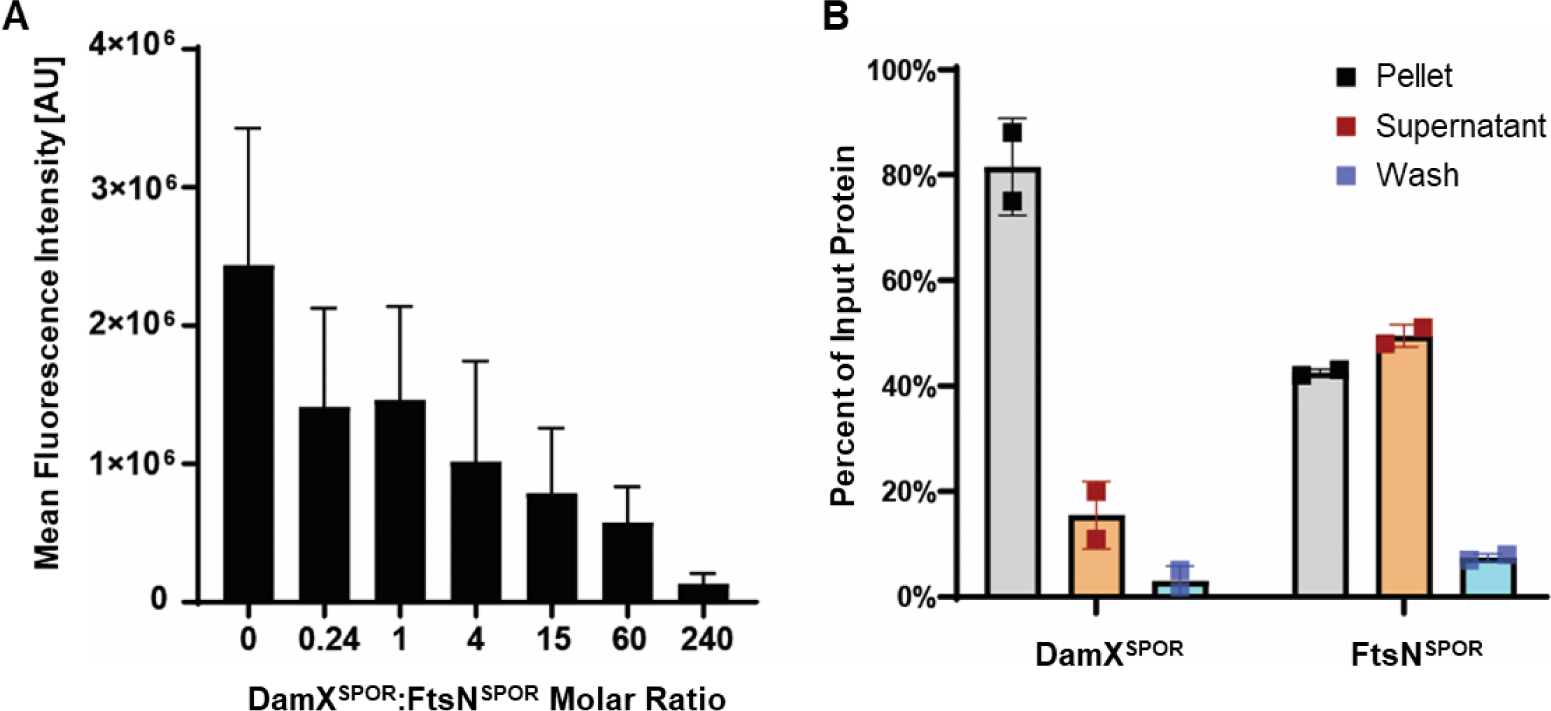
Binding of DamX^SPOR^ and FtsN^SPOR^ to *E. coli* PG sacculi. (A) Competition assay. Mixtures containing 1 µM GFP-FtsN^SPOR^ protein and 0-240 µM non-fluorescent DamX^SPOR^ protein were added to immobilized PG sacculi isolated from a strain lacking five lytic transglycosylases (EC3708). After washing to remove unbound proteins, sacculi were visualized by fluorescence microscopy. Data are the mean ± SE for n ≥ 50 septal bands per protein concentration. AU, arbitrary units. (B) Co-sedimentation assay. DamX^SPOR^ or FtsN^SPOR^ (10 µg) was mixed with sacculi purified from the lytic transglycosylase mutant EC3708. Sacculi were pelleted by ultracentrifugation, washed and centrifuged again. Samples of the supernatant, the wash fluid and the final pellet were subjected to SDS-PAGE and staining with Coomassie to determine the percentage of input protein in each fraction. Data are graphed as the mean ± SEM of n = 2 trials with the value from each trial indicated by a filled square.

We also used a co-sedimentation assay to compare DamX^SPOR^ and FtsN^SPOR^ binding affinity in a semi-quantitative fashion. Mixtures of DamX^SPOR^ or FtsN^SPOR^ with purified *E. coli* sacculi from strain EC3708 were centrifuged to pellet the sacculi along with any bound SPOR protein. The pellets were washed by resuspending the pellets in buffer and then centrifuging again. Samples of the initial supernatant, the wash and the final pellet were analyzed by SDS-PAGE and Coomassie staining. More DamX^SPOR^ than FtsN^SPOR^ co-sedimented with sacculi, consistent with the higher affinity of DamX^SPOR^ determined by ITC. An alternative interpretation is that the fraction of active molecules is higher in the DamX^SPOR^ preparation than in the FtsN^SPOR^ preparation. However, there was more FtsN^SPOR^ than DamX^SPOR^ in the wash fractions, which indicates FtsN^SPOR^ dissociates from PG glycans more readily than does DamX^SPOR^ (**Fig. 3B**).

### Design and Validation of DamX SPOR Domain Swap Constructs

To ask whether the differences in glycan binding affinity observed in ITC are relevant to SPOR domain protein function during cell division, we constructed a set of DamX “swap” proteins with SPOR domains from DedD, FtsN, RlpA or CHU2221 (**Fig. 1B**, **Fig. S2**). In these constructs V377 of DamX is followed by a four amino acid linker (SNNN) and the unique SPOR domain fragments derived from DedD (two constructs, a.a. 141-220 and a.a. 134-220), FtsN (a.a. 240-319), RlpA (a.a. 283-362) and CHU2221 (a.a. 161-249). All constructs were expressed from an IPTG-inducible promoter and included an N-terminal GFP tag. As a positive control, we grafted DamX^SPOR^ (amino acids 338-428) back onto DamX via the same linker sequence used to fuse the other SPOR domains to DamX. The rationale for including two DedD swap constructs with either 0 or 7 amino acids upstream of the start of DedD’s SPOR domain as defined by Uniprot was to see whether such choices affected the outcome of the phenotypic analysis. Negative controls included *gfp* alone and GFP-DamX SPOR domain deletion proteins truncated at V337 or S345 (DamX^ΔSPOR^) (17). The two ΔSPOR constructs behaved identically and were used interchangeably, but figure legends contain strain numbers so the reader can determine which ΔSPOR variant is shown.

All swap construct plasmids and controls were integrated into the chromosome of an *E. coli* Δ*damX* strain so the GFP-DamX^SWAP^ is the only form of DamX in the cells. Swap strains and controls were grown in LB with 10 μM IPTG, which we have previously shown to produce GFP-DamX at physiologically appropriate levels (17). Western blotting revealed that swap constructs and controls were stable and produced at comparable levels to wild-type GFP-DamX (**Fig. 4A**), although the expression level of the CHU2221 construct was consistently about half that of the other swap proteins.

**Figure 4.**
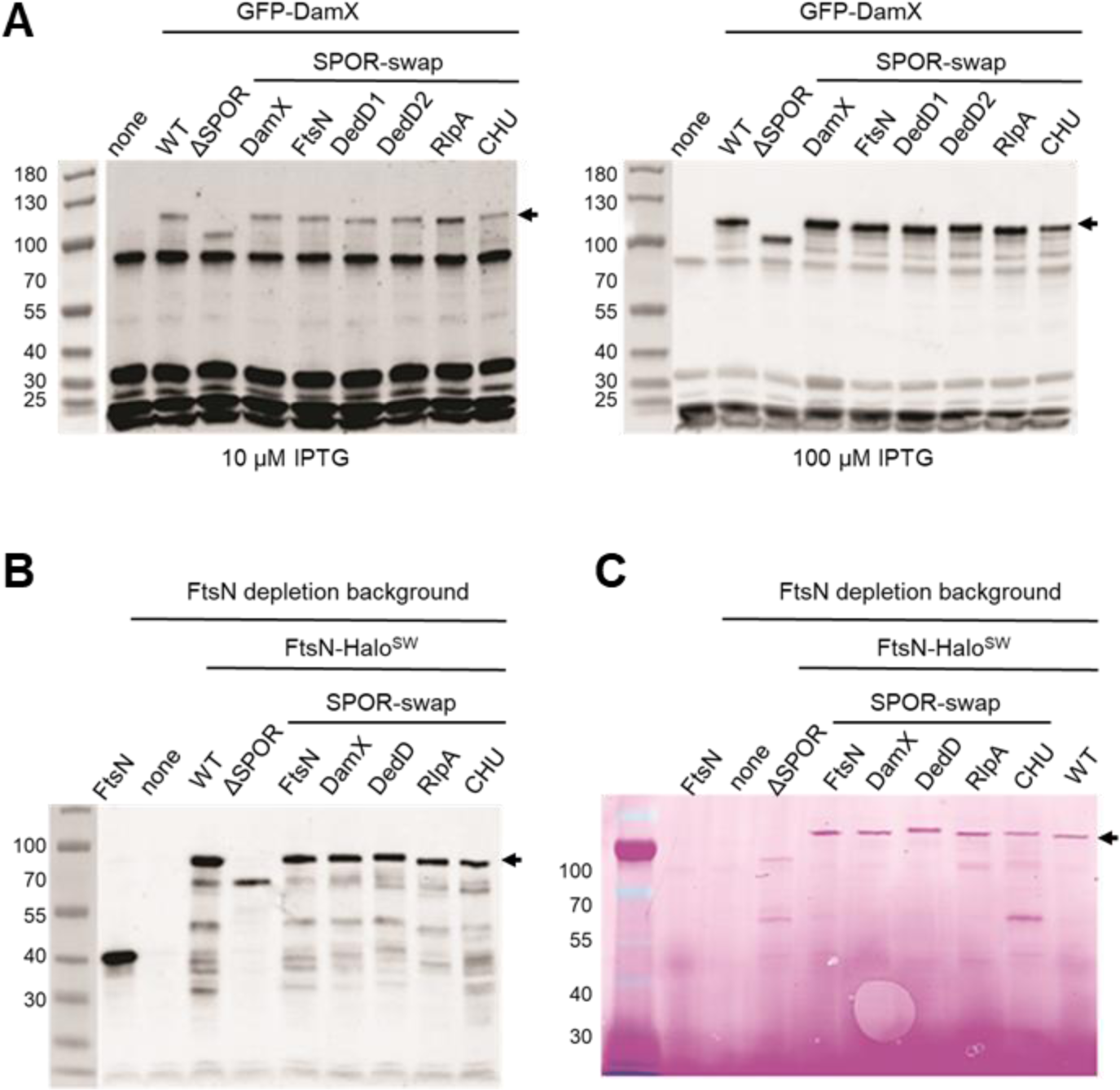
Expression of SPOR domain swap constructs. (A) Analysis of GFP-DamX swap constructs and controls by Western blotting. Whole cell extracts of strains induced with 10 or 100 µM IPTG as indicated were subject to SDS-PAGE, transferred to nitrocellulose and probed with anti-GFP. (B) Analysis of FtsN-Halo^SW^ swap constructs and controls by Western blotting. Whole cell extracts of strains grown in M9-glucose and 20 µM IPTG (or 200 µM IPTG for the CHU SPOR swap) were subjected to SDS-PAGE, transferred to nitrocellulose and probed. (C) Analysis of FtsN-Halo^SW^ swap constructs and controls by Halo ligand binding. Samples grown as in (B) were incubated with JF549 dye, washed, and whole cell extracts were subjected to SDS-PAGE. The gel was imaged in a fluorescence scanner. In all panels molecular mass markers in kilodaltons are indicated to left and arrows point to the swap constructs. Strains in (A) from left to right: EC251, EC2884, EC1910, EC3146, EC3148, EC3150, EC3612, EC3152, and EC3614. Strains in (B) from left to right: EC251, EC1908, EC5676, EC5647, EC5649, EC5651, EC5653, EC5655, and EC5657. Strains in (C) are the same except EC5676 is on the far right.

### DamX SPOR domain swap constructs localize to constriction sites

To evaluate septal localization, cells were grown in lysogeny broth (LB) with 10 µM IPTG to OD600 ∼0.5, fixed and examined by fluorescence microscopy (**Fig. 5A)**. We used fixed cells because septal GFP-DamX increased with time as live cells sat on agarose pads. We first quantified septal localization by scoring cells manually as either positive or negative for a fluorescent band at the midcell. By this criterion, all swap constructs localized in about half of cells in the population, although the high affinity SPOR domain from CHU2221 performed the best (58% septal localization), while the lower affinity SPOR domains from DedD, FtsN and RlpA performed somewhat worse (∼43% septal localization) (**Fig. 5B**). Quantifying septal localization in this way can obscure differences because it fails to distinguish between bright and faint fluorescent bands. To measure septal accumulation of GFP-DamX swap constructs, we quantified the ratio of septal fluorescence to total cellular fluorescence in about 50 cells that exhibited septal localization. By this criterion there was more spread among the swap constructs, with the SPOR domains from CHU2221 and DamX conferring 2- to 3-fold more accumulation of GFP signal at the septum than those from FtsN and RlpA (**Fig. 5B**).

**Figure 5.**
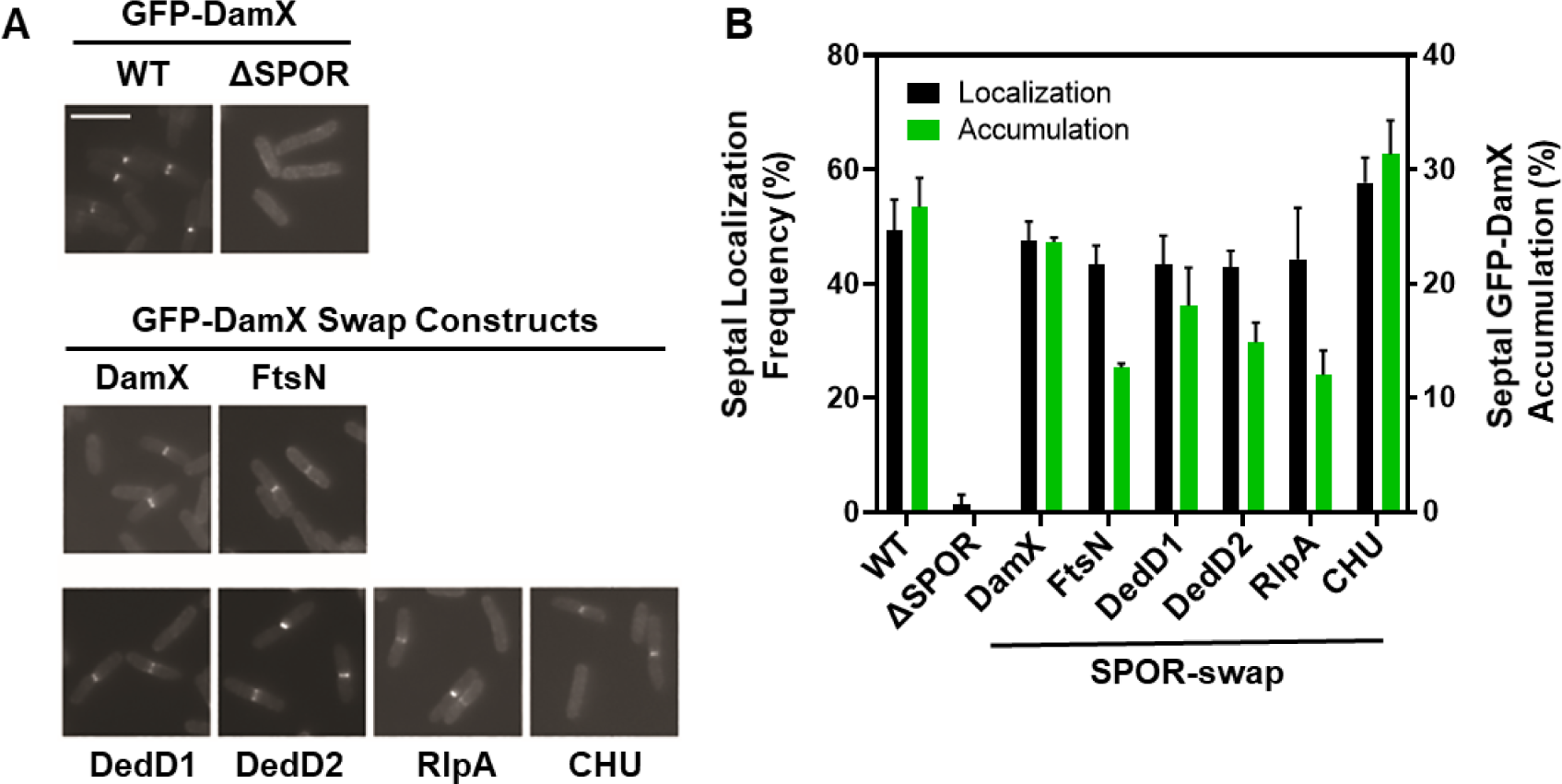
Septal localization of DamX SPOR domain swap constructs. (A) Representative fluorescence micrographs of Δ*damX* strains expressing the indicated *gfp-damX* constructs from an IPTG-inducible promoter integrated into the chromosome at the Φ80 *att* site. Strains were grown in LB with 10 µM IPTG to OD600 = 0.5 and fixed. Size bar = 5 µm. (B) Quantitation of swap construct localization. The fraction of cells exhibiting septal localization was determined manually. Data are graphed as the mean ± SD for three experiments (different days) with ≥ 200 cells scored in each case. Alternatively, septal accumulation of GFP-DamX was determined by dividing the fluorescence at the midcell by the total cell fluorescence. Data are graphed as the mean ± SEM for at least 50 cells that exhibited septal localization. Strains shown are EC2884, EC3146, EC3148, EC3150, EC3612, EC3152, and EC3614.

### DamX SPOR domain swap constructs confer deoxycholate resistance

The absence of DamX renders *E. coli* and *Salmonella* moderately sensitive to the bile salt deoxycholate (9, 16). We confirmed our previous report that expression of a wild-type *gfp-damX* fusion with 10 µM IPTG restores full deoxycholate resistance to a Δ*damX* mutant, but a *gfp-damX*^ΔSPOR^ allele does not (17) (**Fig. 6A**). Interestingly, all of our GFP-DamX^swap^ proteins restored deoxycholate resistance, although the lowest affinity SPOR domain from FtsN was less effective than the others (**Fig. 6A**).

**Figure 6.**
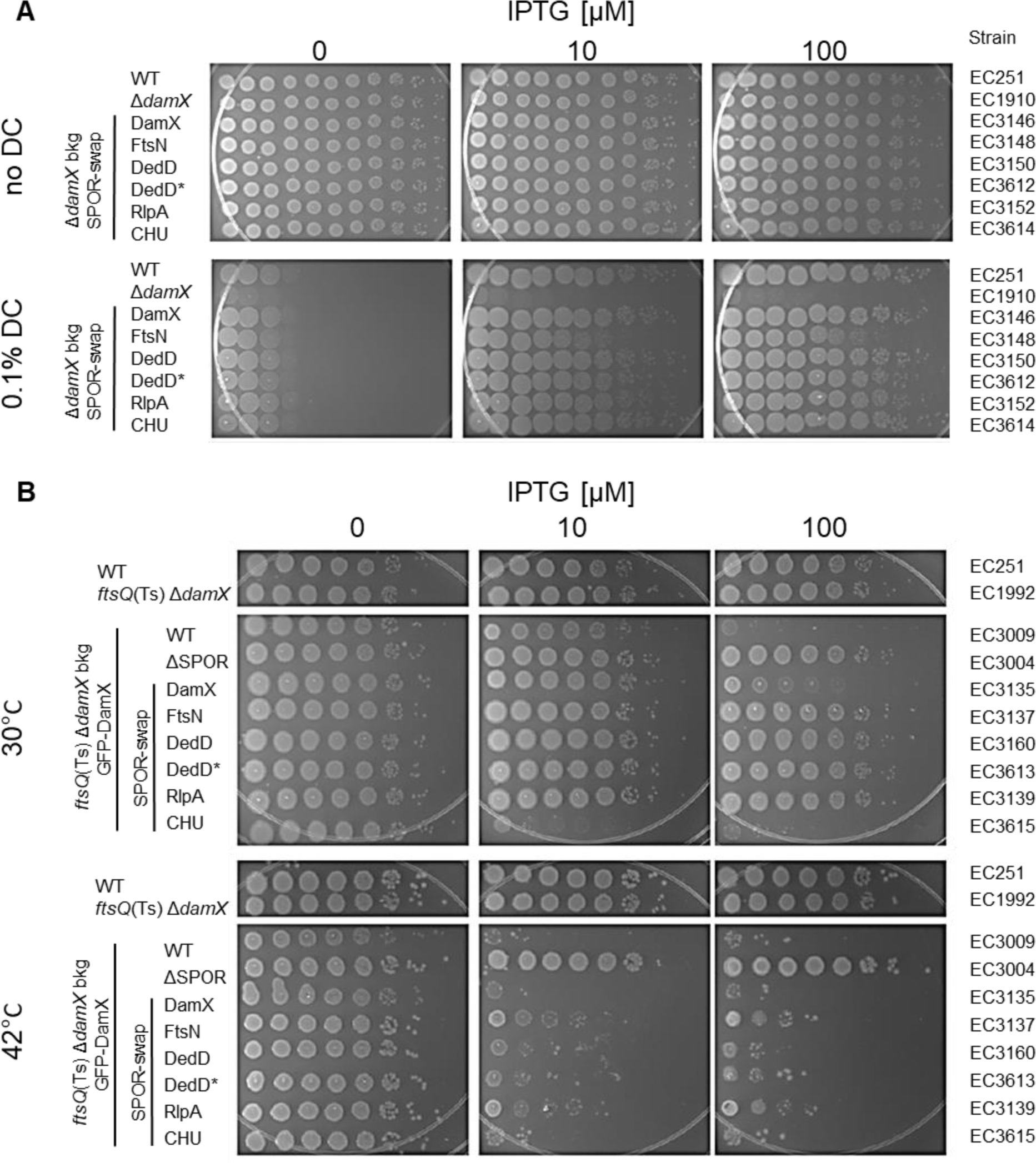
Spot titer assays of DamX swap construction functionality. (A) Deoxycholate (DC) resistance of EC1910 [Δ*damX*] derivatives expressing the indicated *gfp-damX* constructs under IPTG control. Overnight cultures grown at 30°C in LB were adjusted to OD_600_ = 1.0 in LB0N. Then 4-fold serial dilutions were prepared in LB0N and spotted onto LB0N plates with or without DC and IPTG as indicated. Plates were incubated at 30°C for 18 hr. The wild-type (i.e., *damX* ) parent EC251 is included for comparison. (B) The Ts phenotype of EC1992 [*ftsQ1*(Ts) Δ*damX*] derivatives expressing the indicated *gfp-damX* constructs under IPTG control. Overnight cultures grown at 30°C in LB were transferred to LB0N, serially diluted 10-fold, and spotted onto LB0N plates with IPTG as indicated. Plates were incubated at 30°C or 42°C for 16 hr. The wild-type (i.e., *damX*^+^) parent EC251 is included for comparison.

### DamX SPOR domain swap constructs render *ftsQ1*(Ts) lethal at 42°C

We have shown previously that deleting *damX* from an *ftsQ1*(Ts) strain restores viability at the normally non-permissive temperature of 42°C (9). The underlying mechanism is unknown, but overproduction of FtsN also rescues *ftsQ1*(Ts) (48), so the absence of *damX* might increase FtsN localization. Expressing *gfp-damX* at physiological levels restores the Ts phenotype, whereas comparable expression of *gfp-damX*^ΔSPOR^ does not (**Fig. 6B**) (17). All of our swap constructs also restored the *ftsQ1*(Ts) lethal phenotype when induced with 10 µM IPTG (**Fig. 6B**). Interestingly, overproducing GFP-DamX by inducing with 100 µM IPTG prevented the *ftsQ1*(Ts) mutant from forming a colony even when plated at 30°C (**Fig. 6B**). A swap construct with the tighter binding SPOR domain from CHU2221 prevented growth even when induced with only 10 µM IPTG (**Fig. 6B**). Curiously, a swap construct in which DamX^SPOR^ was grafted via a linker back onto DamX was not inhibitory, indicating this swap is not completely functional.

This finding is consistent with the somewhat reduced septal localization of the GFP-DamX SPOR domain self-swap compared to the wild-type GFP-DamX fusion protein (**Fig. 5B**).

### DamX SPOR domain swap constructs differ in their ability to promote and inhibit cell division

A simple Δ*damX* null mutant has no obvious division defect and cell length is essentially wild-type (8, 9), so it is not feasible to use rescue of a Δ*damX* null mutant to test whether the swap constructs function normally in cell division. Instead we integrated our swap plasmids and controls into a Δ*damX* Δ*dedD* double mutant, which has a synthetic division defect and averages 10-15 µm in length in different studies (8, 9, 17). We have shown previously that expressing *gfp-damX* at physiologically appropriate levels in the Δ*damX* Δ*dedD* background improves division and cells become shorter (17). Conversely, overexpressing *gfp-damX* inhibits division and cells become longer (17). Here we observed that the Δ*damX* Δ*dedD* double mutant had a cell length of ∼13 µm, which decreased to ∼ 9 µm when *gfp-damX* was induced with 10 µM IPTG and increased to ∼100 µm at 100 µM IPTG (**Fig. 7A, B**). The DamX SPOR domain self-swap control behaved similarly in this assay, as did a swap construct with the high-affinity SPOR domain from CHU2221. In contrast, none of the swap constructs with low-affinity SPOR domains (DedD^SPOR^, FtsN^SPOR^, or RlpA^SPOR^) either rescued or inhibited cell division (**Fig. 7B, S3**), even though they localized sharply to potential division sites in the double mutant background (**Fig. S4**).

**Figure 7.**
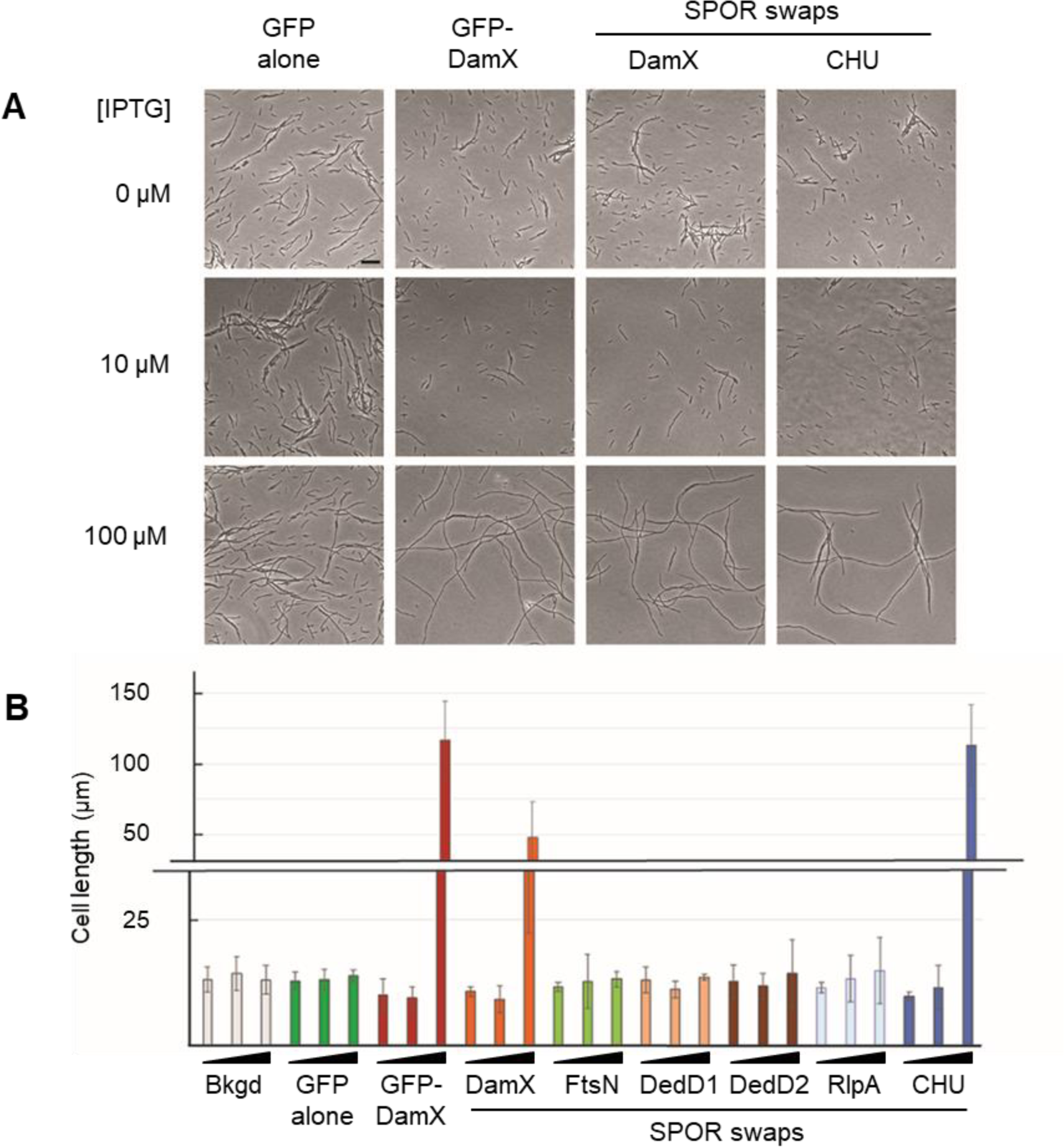
Ability of DamX SPOR swap constructs to improve or inhibit cell division. (A) Phase contrast micrographs of EC1926 [Δ*dedD* Δ*damX<>kan*] derivatives producing the indicated constructs from an IPTG-inducible promoter. Size bar = 10 µm. (B) Cell lengths. Wedges indicate cultures induced with 0, 10 or 100 µM IPTG. Data are graphed as the geometric mean ± SD from 3-5 experiments with ≥ 100 cells (or filaments) measured per experiment. Strains in (A) from the left: EC2315, EC2316, EC3092, and EC3608. Strains in (B) from the left: EC1926, EC2315, EC2316, EC3092, EC3098, EC3094, EC3607, EC3142, and EC3698.

### Design of FtsN SPOR Domain Swap Constructs

FtsN SPOR domain swap constructs were derived from a functional FtsN-Halo sandwich fusion (FtsN-Halo^SW^) in which a HaloTag is inserted between E60 and E61 of FtsN (**Fig. 1B**) (49). This insertion site places Halo in the periplasm, shortly downstream of FtsN’s transmembrane helix but before the essential domain (^E^FtsN) required to activate constriction (8, 50). The FtsN SPOR domain swap constructs and controls were integrated into the chromosome and expressed from an IPTG-inducible promoter in a PBAD::*ftsN* depletion strain. Western blotting with anti-FtsN sera verified swap construct expression and effective depletion of native FtsN when strains were grown in M9-glucose containing IPTG (**Fig. 4B**). About 10-fold more IPTG was needed to induce the CHU2221 swap to match its abundance to the other swap proteins. Because our anti-FtsN sera was raised against the entire periplasmic domain of FtsN, swap constructs are less cross reactive than constructs that retain FtsN’s native SPOR domain. For this reason we also compared swap construct expression levels using the HaloTag ligand JF549 to label the proteins (**Fig. 4C**). Note that while the JF549 dye allows for comparison of the various Halo^SW^ constructs, it cannot be used to benchmark expression to native FtsN in wild-type *E. coli*.

### FtsN SPOR domain swap constructs localize to constriction sites and support cell division

To assess swap construct functionality, cultures were grown at room temperature in M9-glucose containing IPTG to OD600 ∼0.5, labeled with the HaloTag ligand JF549, then fixed, and examined by phase contrast and fluorescence microscopy. Cells were scored manually for the presence or absence of a fluorescent band at the midcell (**Fig. S5**, **Table 2**). A wild-type FtsN-Halo^SW^ construct localized to division sites in about 20% of cells. As expected, septal localization decreased to near zero when the SPOR domain was deleted (8). The small amount of residual localization in the absence of a SPOR domain can be explained by FtsN’s interactions with other division proteins in the cytoplasm and the periplasm (8, 11, 51–53).

**Table 2.**
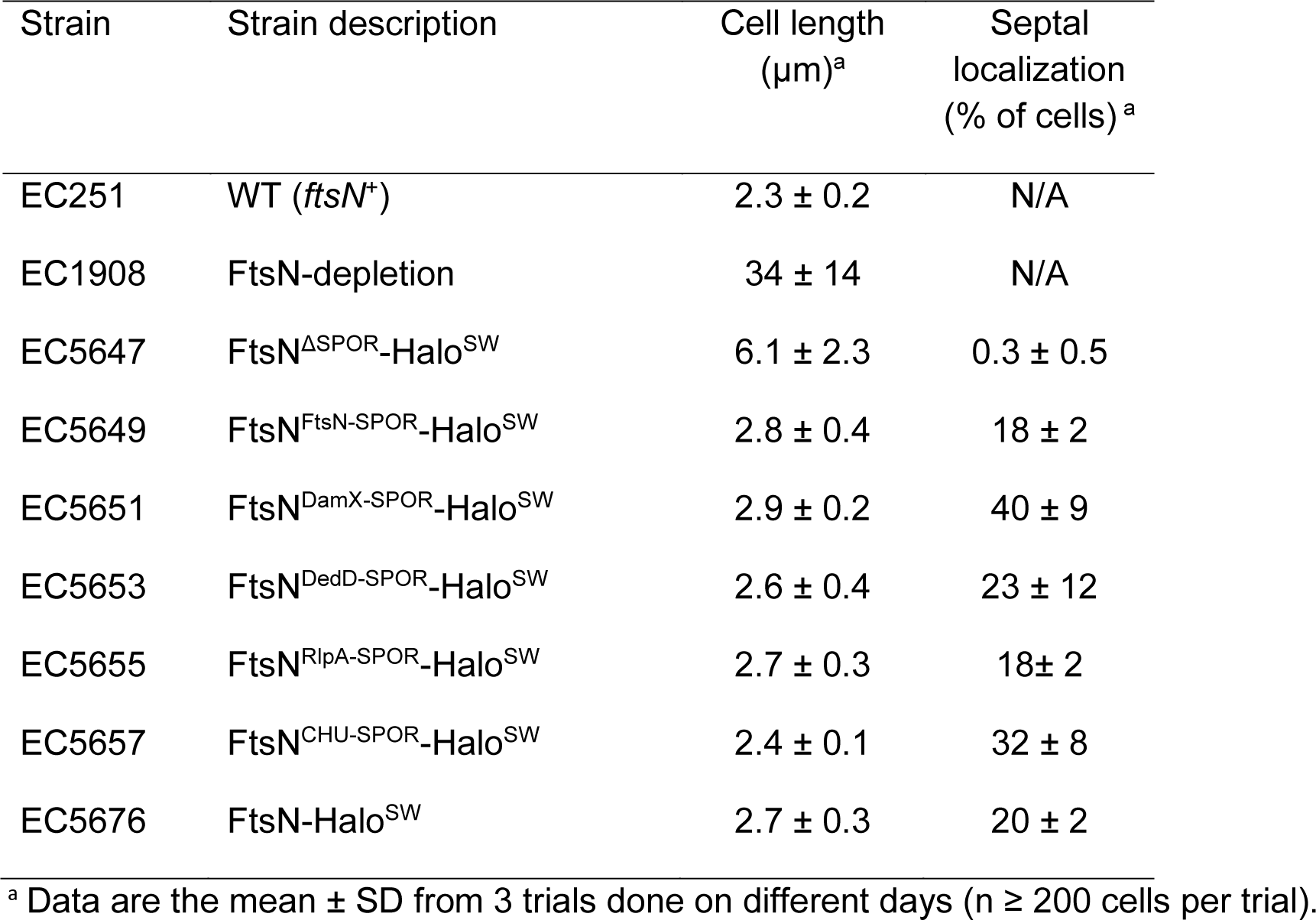
Phenotypic characterization of FtsN-Halo^SW^ SPOR domain swap strains

Normal frequencies of septal localization were restored by adding back the SPOR domain from FtsN (as a control), DedD, or RlpA, while the higher affinity SPOR domains from DamX or CHU2221 increased localization frequency to ∼30% or 40% of the cells (**Fig. 6**).

We assessed the functionality of FtsN SPOR domain swap constructs in cell division by measuring cell length (**Fig. S5**, **Table 2**). When grown in M9-glucose, our lab MG1655 isolate (EC251) averaged 2.3 µm in length. Depletion of FtsN impaired constriction, so average cell length increased to about 40 µm. Producing FtsN-Halo^ΔSPOR^ in the depletion background largely corrected the division defect, although cells were still about twice as long as wild-type (∼5 µm). This result was expected because it is well-established that FtsN’s SPOR domain improves cell division but is not required for it (8, 10, 14). Cells of the depletion strain that expressed wild-type FtsN-Halo^SW^ or any of the corresponding FtsN-Halo^SW^ SPOR domain swap derivatives were close to normal length (range 2.3-2.9 µm). Thus, all of the FtsN SPOR domain swap constructs function well in *E. coli* cell division.

## Discussion

SPOR domains are small, peptidoglycan (PG) binding domains that target proteins to the division site by binding to the glycan backbone of PG at sites where cell wall amidases have removed stem peptides (8, 14, 36, 37). These PG sites, referred to here as denuded glycans, are transiently enriched in septal PG by cell wall amidases involved in daughter cell separation (27, 29–31). If all SPOR domains bind to the same site, they might be functionally interchangeable. In support of that notion, it has previously been shown that a chimeric *C. crescentus* FtsN protein equipped with the SPOR domain from FtsN of *E. coli* supported cell division in *C. crescentus* (10). But evidence for differences in SPOR domain functionality emerged from a study showing differential localization frequencies in *E. coli* cells treated with cephalexin to inhibit synthesis of septal PG (8). Moreover, SPOR domains are not well conserved at the level of amino acid sequence (9, 17), which is suggestive of functional differences.

Here we used a combination of *K*d determination by ITC and domain swapping to explore these ideas. The SPOR domains we chose to work with came from four *E. coli* division proteins (DamX, DedD, FtsN and RlpA) and one uncharacterized *Cytophaga hutchinsonii* protein (CHU2221). This set of SPOR domains exhibits <20% identity in all possible pairwise comparisons (9). We found that binding affinities of these SPOR domains were all in the low micromolar range but differed by up to about 10-fold: CHU2221^SPOR^ 0.8 µM ≈ DamX^SPOR^ 1.3 µM < DedD^SPOR^ 3.7 µM ≈ RlpA^SPOR^ 5.1 µM < FtsN^SPOR^ ∼10.2 µM (**Fig. 2**, **Fig. S1**, **Table 1**). These findings were confirmed qualitatively by showing DamX^SPOR^ binds PG more tightly than FtsN^SPOR^ in a co-sedimentation assay (**Fig. 3B**).

Although most of our ITC measurements employed denuded glycans from *S. aureus* PG, we think they are applicable to denuded glycans from *E. coli* PG because we have shown previously that DamX^SPOR^ localizes to septal regions of sacculi from both organisms (1, 36).

Moreover, we found here by ITC that DamX^SPOR^ binds denuded glycans from both organisms with very similar affinities of 1.7 and 1.3 µM for *E. coli* and *S. aureus*, respectively (**Fig. 2**, **Fig. S1**, **Table 1**). These findings make sense because *E. coli* and *S. aureus* PG differ primarily with respect to the structure of the stem peptides and crosslinks (23), neither of which are relevant to SPOR domain binding (36, 37, 54). There is, however, a noteworthy difference in the glycan backbone—the C6 hydroxyl of NAM is often acetylated in *S. aureus* but not *E. coli* (23, 54, 55).

However, it is unlikely that the C6 hydroxyl of NAM retained any O-acetyl groups because they should be lost during the acid incubation step used to remove teichoic acids during isolation of *S. aureus* sacculi. In addition, the C6 hydroxyl of NAM is solvent exposed in the crystal structure of a *Pa*RlpA^SPOR^-glycan complex (37), implying SPOR domain binding would not be affected even if there are residual C6 O-acetyl groups in our glycan preparations.

SPOR domain swapping was used to assess the potential biological significance of the different glycan binding affinities of the various SPOR domains. As host proteins for SPOR domain replacement, we selected FtsN and DamX of *E. coli*. Both of these proteins are involved in cell division, with FtsN being essential and DamX being non-essential (8, 9, 48). Previous reports had shown that FtsN function in cell division is only modestly impaired by removal of its SPOR domain (8, 10, 14), limiting the range of phenotypic differences expected from domain swapping. Conversely, DamX functions are highly dependent upon its SPOR domain (17).

As expected, deleting the SPOR domain from FtsN of *E. coli* nearly eliminated septal localization while impairing cell division only modestly (**Table 2**, **Fig. S5**). Adding back any of four other SPOR domains (DamX, DedD, RlpA, CHU2221) restored septal localization and conferred near normal function in cell division as reflected in average cell length (**Table 2**, **Fig. S5**). Thus, FtsN functions reasonably well with a variety of SPOR domains, including SPOR domains with a 10-fold greater affinity for denuded glycans. FtsN was recently discovered to move processively around the septum at an average velocity ∼9 nm s^-1^ as part of the FtsWI complexes that are actively synthesizing sPG (49). The average run length for FtsN in these complexes is ∼150 nm (49), over twice the ∼70 nm length of FtsN’s periplasmic linker domain (41), so we think FtsN’s SPOR domain must let go of denuded glycans to promote FtsWI activity. Thus, it is intriguing that the tighter-binding SPOR domains from DamX and CHU2221 did not prevent FtsN from supporting cell division.

Our results with DamX were more nuanced. Unlike FtsN, DamX requires its SPOR domain for septal localization and function in a variety of assays (17). Here we observed that DamX swap proteins containing any of four heterologous SPOR domains localized to the midcell and functioned like authentic DamX when it came to conferring normal deoxycholate resistance or the canonical *ftsQ*(Ts) phenotype (**Fig. 5, 6**). But differences became apparent when the swap proteins were tested for their ability to improve cell division when produced at normal levels in a *ΔdamXΔdedD* double mutant or to inhibit cell division when overproduced in that same background (**Fig. 7**, **Fig. S3, S4**). In these assays, only the high-affinity *Cytophaga* SPOR domain from CHU2221 functioned like authentic DamX^SPOR^. The failure of the DedD, FtsN and RlpA SPOR domain swaps in these assays is remarkable in light of the fact that these constructs accumulated well at potential division sites. The simplest interpretation of these results is that DamX function requires not only midcell localization but also strong anchoring to denuded glycans in septal PG. This might indicate DamX plays a structural role in tethering the PG layer to the cytoplasmic membrane during constriction, and could be related to why DamX is about 4-fold more abundant than the other three *E. coli* SPOR domain proteins (56). It will be interesting to learn if septal DamX is less dynamic than FtsN.

Although all four *E. coli* SPOR domains localize strongly to constriction sites, DedD^SPOR^ and RlpA^SPOR^ also localize weakly to the cell cylinder, suggesting these SPOR domains might have somewhat different binding-site requirements than do DamX^SPOR^ and FtsN^SPOR^ (8, 9, 36).

Nevertheless, we think that differences in glycan binding affinity rather than differences in binding site preferences best account for the results of our SPOR domain swapping experiments. (i) All four *E. coli* SPOR domains clearly bind denuded glycans, as evidenced by dramatically increased binding to amidase-treated PG and dramatically decreased binding to PG from which denuded glycans have been removed by digestion with a lytic transglycosylase (36). (ii) DamX^SPOR^ and FtsN^SPOR^ compete for binding to septal PG in isolated sacculi, demonstrating directly that they utilize the same sites. (**Fig. 3A**). (iii) In a previous study, we characterized a DamX mutant with a W416L substitution in the SPOR domain glycan-binding site. The W416L substitution caused modest decreases in PG-binding, septal localization and deoxycholate resistance but abolished the ability of DamX to promote or inhibit division in a *ΔdamXΔdedD* double mutant background (17). Thus, the consequences of weakening DamX’s SPOR-glycan interaction with a W416L lesion are similar to those of replacing DamX’s high affinity SPOR domain with the lower affinity SPOR domains from DedD, FtsN or RlpA. (iv) Docking models derived from the *Pa*RlpA^SPOR^-glycan structure predict the SPOR domains from DamX, DedD, FtsN and RlpA all bind to a NAG-NAM-NAG-NAM tetrasaccharide (37).

In summary, bacterial cell division requires remodeling the cell envelope at the right place and the right time. SPOR domains contribute to this process by delivering division proteins to constriction sites in response to the appearance of denuded PG glycans (1). If septal targeting were the only function of SPOR domains, they ought to be fully interchangeable. This appears be the case for FtsN but not DamX, which requires a SPOR domain with high glycan-binding affinity to function well in cell division. Further work will be needed to determine how glycan-binding affinity contributes to DamX function. One possibility is that affinity differences determine priority when multiple SPOR domain proteins are competing for denuded glycans.

Alternatively, the on- and off-rates for PG binding might be important for SPOR protein dynamics at the septum.

## Materials and Methods

### Media

Lysogeny broth (LB) contained 10 g tryptone, 5 g yeast extract and 10 g NaCl per liter, and was solidified with 15 g agar for plates. NaCl was omitted when making LB0N. Tryptic Soy Broth (Bacto) was used to grow *S. aureus* for sacculi preparations. For FtsN-Halo^SW^ experiments, cells were grown in M9-glucose minimal medium (0.4% D-glucose, 1× MEM amino acids and 1× MEM vitamin) prepared with ingredients from Sigma (57). Antibiotics were used at the following concentrations: ampicillin, 200 µg/ml; chloramphenicol, 20 µg/ml; kanamycin, 40 µg/ml. Genes under PBAD control were induced with 0.2% L-arabinose (58). Genes under control of modified *Trc* promoters were induced with Isopropyl-β-D-thiogalactoside (IPTG) as indicated (59).

### Bacterial strains, plasmids, and oligonucleotide primers

Bacterial strains, plasmids and oligonucleotides are listed in Tables S1, S2 and S3 and described in the supplemental materials. Strain construction was by transformation or CRIM plasmid integration and followed published procedures (60, 61). Plasmid construction used restriction enzymes, T4 DNA ligase, Q5 DNA polymerase and NEBuilder HiFi DNA Assembly Master Mix from New England Biolabs (NEB). Plasmid DNA was purified using the QIAprep Spin Miniprep Kit (Qiagen). DNA fragments were recovered from PCR reaction mixtures and agarose gels using Monarch kits (NEB). Oligonucleotides were obtained from Integrated DNA Technologies.

### Microscopy and image analysis of *gfp-damX* swap constructs

To determine localization efficiency of *gfp-damX^swap^* constructs, overnight cultures grown in LB at 30°C were diluted 1:500 into LB containing 10 µM IPTG and grown to OD600 = 0.5. Cells were fixed with 2.5% paraformaldehyde in 20 mM NaPO4 buffer, pH 7.4, as described (20). Cells were immobilized on 1% agarose pads for imaging. Phase-contrast and fluorescence micrographs were captured with an Olympus BX60 microscope equipped with a Hamamatsu ORCA-Flash4.0 V2+ complementary metal-oxide semiconductor (CMOS) camera, Olympus 100X UPlanApo objective (numerical aperture 1.35), Olympus 100W High Pressure Mercury Burner (BH2-RFL- T3), and Olympus cellSens Dimension acquisition software. Green fluorescence was visualized with a GFP filter set (catalog no. 49002, Chroma Technology Corp.). Typical exposure times for GFP were 1 s. Cells were scored manually as positive or negative for septal localization.

Fractional accumulation of GFP-SPOR at the midcell was determined in ImageJ by dividing the fluorescence at the midcell by the total cell fluorescence. Adobe Illustrator or Olympus cellSens Dimension software were used to adjust image brightness and contrast, to crop images, and to assemble images into figures.

To determine cell morphology and localization of *gfp-damX^swap^* constructs in a Δ*damX* Δ*dedD* background, cultures were processed similarly. Cell length measurements were made using ImageJ or Olympus cellSens Dimension software. Olympus 40X UPlanFI objective lens (numerical aperture 0.75) was used for capturing images for cell division inhibition experiment.

### Microscopy of FtsN-Halo^SW^ constructs labeled with JF549 dye

FtsN-Halo^SW^ swap strains were grown at room temperature in M9-glucose supplemented with amino acids, vitamins and IPTG as described (49). When cultures reached OD600 ∼ 0.35, a 100 µl aliquot was removed and incubated with 1 µM JF549 HaloTag ligand in the dark for 30 min. Cells were then fixed with paraformaldehyde as described above. Microscopy was performed as above except that in this case red fluorescence was visualized with a Texas Red filter set (catalog no. 41004, Chroma Technology Corp.). Typical exposure time was 400 msec.

### Western blotting

In Western blotting of GFP-DamX swap constructs, 1.0 ml of culture at an OD600 of 0.50 was centrifuged and the cell pellet taken up in 100 µl of 1x SDS-PAGE loading buffer to achieve a sample concentration of 5.0 OD600 units/ml. For FtsN-Halo^SW^ constructs, cultures were harvested at OD600 = 0.35 and the harvest volume was increased to 1.4 ml. Samples were heated 10 min at 95°C before loading 10 µl aliquots onto precast mini-PROTEAN TGX gels (Bio-Rad). The polyacrylamide concentration was 7.5% for GFP-DamX constructs, 10% for FtsN-Halo^SW^ constructs. Electrophoresis, transfer to nitrocellulose, and blot development followed standard procedures. Blocking was with 5% nonfat dry milk diluted in phosphate-buffered saline containing 0.1% TWEEN 20 (PBST). Primary antibody for GFP-DamX constructs was polyclonal rabbit anti-GFP (ThermoFisher) serum diluted 1:10,000 in PBST. For FtsN-Halo^SW^ constructs it was anti-FtsN sera that had been pre-absorbed against a lysate of DH5α/pMAL-C2 and then used at a 1:1000 dilution. Secondary antibody was horseradish peroxidase-conjugated goat anti-rabbit antibody (1:8,000) (Pierce), which in turn was detected with SuperSignal Pico West PLUS chemiluminescent substrate (Thermo Scientific). Blots were visualized with a ChemDoc Touch imaging system (Bio-Rad).

### Detection of JF549-labeled FtsN-Halo^SW^ swap constructs after SDS-PAGE

As an alternative to Western blotting, the expression and stability of FtsN-Halo^SW^ swap constructs was assessed using JF549 dye as follows. One ml of culture grown for microscopy was incubated with 1 µM JF549 dye for 30 min at room temperature in the dark. Cells were pelleted and taken up in ∼ 70 µl 1x SDS-PAGE loading buffer to achieve a sample concentration of 5.0 OD600 units per ml. Samples were heated for 10 min at 95°C before loading 10 µl onto a 10% polyacrylamide precast mini-PROTEAN TGX gel (Bio-Rad). After electrophoresis the gel was washed twice with distilled water, then imaged on a Sapphire Biomolecular Imager (Azure Biosystems) using the Alexa 546 setting (excitation 520 nm, emission 565 nm) with scanning parameters set to scan gel, 100-pixel resolution, and slow speed.

### Effect of GFP-DamX^swap^ proteins on the *ftsQ1*(Ts) phenotype

*E. coli ftsQ1*(Ts) Δ*damX*<>*kan* strains harboring P204::*gfp-damX^swap^* constructs (or controls) integrated at the Φ80 *attP* site were grown overnight in LB media with 40 µg/ml kanamycin. The next day, 200 µl of culture was centrifuged, and the resulting cell pellet was taken up in ∼1 ml of LB0N to achieve an OD600 = 1.0. A series of 10-fold serial dilutions in LB0N medium was prepared, and 4 µl of each dilution were spotted onto duplicate LB0N plates containing 0, 25, or 100 µM IPTG as indicated. Plates were incubated at 30°C or 42°C for 16 hours, stored at 4 °C for a day, and then photographed.

### Effect of GFP-DamX^swap^ proteins on deoxycholate sensitivity

*E. coli* Δ*damX*<>*kan* strains harboring P204::*gfp-damX^swap^* constructs (or controls) integrated at the Φ80 *attP* site were grown overnight in LB media with 40 µg/ml kanamycin. The next day, cultures were adjusted to an OD600 of 1.0 in LB0N medium. A series of 4-fold serial dilutions were prepared and 3 µl of each dilution were spotted onto LB0N plates with and without 0.1 % deoxycholate and 0, 10, or 100 µM IPTG as indicated. Plates were incubated at 30°C for 18 hours, stored at 4 °C for a day, and then photographed.

### Protein Purification

All proteins were overproduced in *E. coli* BL21(DE3) derivatives except for His6-FtsN^SPOR^ and His6-GFP-FtsN^SPOR^, which were overproduced in SHuffle T7 in order to promote formation of disulfide bonds in FtsN’s SPOR domain (42, 62). All purification steps were conducted at 4°C and proteins were purified using Talon cobalt affinity resin (Clonetech) as described (36). Purified His6-SPOR proteins to be used for ITC measurements were dialyzed against ITC sample buffer (25 mM NaPO4, pH 7.5, 150 mM NaCl) and stored at 4°C until use, typically within 2 weeks. For competition and co-sedimentation assays (Fig. 2), His6-DamX^SPOR^, His6-FtsN^SPOR^ and His6-GFP-FtsN^SPOR^ were purified by the same protocol but dialyzed against storage buffer (50 mM Tris-HCl, pH 7.5, 200 mM NaCl, 5 % Glycerol), and aliquots were stored at -80°C until needed. His6-AmiD and His6-Atl amidase were purified and stored similarly. In all cases, protein purity was ∼95% as judged by SDS-PAGE. The concentrations of purified proteins were determined by absorbance at 280 nm using an extinction coefficient based on the amino acid sequence of each protein construct. Typical yields per liter of culture were 3 to 5 mg, except that for FtsN proteins, which require formation of an essential disulfide bond, the yield was 0.2 to 0.6 mg per liter of culture.

### Preparation of *E. coli* sacculi

*E. coli* sacculi were isolated from the multiple lytic transglycosylase mutant E3708 as described (36), starting from 2 liters of culture (4 x 500 ml in 2 L flasks) at an OD600 = 0.5. Isolated sacculi were suspended in a final volume of 450 µl of nanopure H2O. Assuming no losses along the way, these suspensions contained about 4.4 x 10^8^ sacculi per µl. We used 200 µl of the suspension per titration assay (∼8.8 x 10^10^ sacculi). To obtain AmiD-treated sacculi for ITC, sacculi from 2 liters of culture were incubated with 100 µg AmiD per ml for 2 hr at 37°C prior to α-chymotrypsin treatment. At the end of the isolation procedure, AmiD-treated sacculi were suspended in 450 µl Nanopure water and 100-200 µl of the suspension was used for one titration assay (4.4 to 8.8 x 10^10^ sacculi).

### Preparation of soluble denuded PG glycans from *S. aureus*

Sacculi were isolated from 1 liter cultures of *S. aureus* grown in Bacto Tryptic Soy Broth at 30°C to OD600 = 0.5. The procedure for isolating sacculi was as described for *B. subtilis* (36, 63). Purified sacculi were incubated with 100 µg/ml Atl amidase for 48 hours at 37°C in 50 mM Tris-HCl pH 7.0 to remove stem peptides. Smaller PG hydrolysis products were separated from large PG fragments and Atl amidase by ultrafiltration using a 30 kDa device (Amicon catalog #UFC9030), saving the filtrate. The filtrate was then applied to a 3 kDa device (Amicon catalog #UFC9003) to remove free peptides, this time saving the retained fraction. Buffer components and small PG degradation fragments were removed by adding 2 ml of H2O to the retentate and recentrifuging for a total of at least 7 exchanges against water. The retained fraction was lyophilized and stored at -80°C. The isolated glycans are a white powder with a needle-like shape. Yields range from 1 to 5 mg from a liter of cells.

### Isothermal titration calorimetry (ITC)

All experiments were conducted on a GE MicroCal VP-ITC System (GE Healthcare) at 25 °C. All titrations were performed in 25 mM NaPO4, pH 7.5, 150 mM NaCl. The SPOR domain proteins and ligands were degassed. SPOR proteins were loaded in the syringe as the injected sample, while NAG, chitosan, sacculi, or glycans were placed in the sample cell. SPOR protein injections of 14 μl over 20 total injections were added to the sample cell containing 1.4 ml of glycan or other test ligand. An injection duration time of 14 s and a spacing of 240 s were set for each injection. Injection of SPOR protein into buffer showed constant heats of dilution. Heats of dilution were determined by averaging the heat evolved by the last five injections and subtracted from the raw data. The values for affinity were then determined using the Origin software provided by the manufacturer. Replicates for each run were further analyzed using GraphPad Prism 7. The concentration of SPOR domain was based on A280, the calculated molecular mass of each His6-tagged domain, the assumption that preparations were 100% active for binding and the assumption (based on a published structure) that SPOR domains bind as monomers to an octasaccharide glycan site (although the direct contacts are to a tetrasaccharide). The concentration of binding sites in AmiD-treated *E. coli* sacculi was set arbitrarily to 20 µM. The concentration of glycan binding sites in soluble denuded *S. aureus* glycans was calculated based on the following information: glycan dry weight, the molecular mass of a NAG-NAM disaccharide (479 g/mol), and 4 NAG-NAM disaccharides per binding site (i.e., an octasaccharide). Muropeptide analysis indicated <10% of NAM moieties retained a stem peptide, so the contribution of stem peptides to glycan dry weight was ignored. The amounts of SPOR domain protein and ligand varied across our ITC experiments. SPOR domains were loaded in the syringe at 50-250 µM. Ligands were added to the sample cell at the following concentrations: denuded glycan, 2 to 50 µM estimated binding site; NAG, 20 µM; soluble chitosan MW 5000, 20 µM. NAG and chitosan were from Sigma (catalog numbers A8625 and 523682, respectively).

### Competition assay

The assay was adapted from (36). His6-GFP-FtsN^SPOR^ (1 µM) was mixed with increasing amounts (0-240 molar ratio) of His6-DamX^SPOR^ protein in phosphate buffered saline (PBS: 137 mM NaCl, 3 mM KCl, 9 mM, NaH2PO4, and 2 mM KH2PO4, pH 7.4) containing bovine serum albumin (BSA) at 20 mg/ml. Protein mixtures were added to sacculi from strain EC3708 that had been immobilized on a poly-L-lysine-treated glass slide. After incubation for 30 min to allow SPOR domain proteins to bind, sacculi were washed 5 times with PBS then visualized by fluorescence microscopy. Septal localization of GFP-FtsN^SPOR^ was quantified as the ratio of septal fluorescence to the fluorescence of the entire sacculus.

### Co-sedimentation assay

Co-sedimentation assays were conducted in 25 mM NaPO4 (pH 7.5) and 150 mM NaCl (42). Binding mixtures for the co-sedimentation assay contained 10 µg of SPOR domain protein and 20 µl of a suspension of PG sacculi from EC3708 (∼9 x 10^9^ sacculi) in a final volume of 100 µl. After incubation at 4°C for 1 hour, mixtures were centrifuged at 4°C in a Beckman TLA-55 rotor at 112,000 x g (average) for 45 min. The supernatant containing non-bound SPOR protein was saved for analysis. The pellet was washed by suspending in 100 µl of cold assay buffer and centrifuging as above. The wash supernatant was saved for analysis. The pellet was suspended again in 100 µl and saved for analysis. All fractions (supernatant, wash, pellet) were subjected to SDS-PAGE and Coomassie staining.

Gels were analyzed using a ChemiDoc Touch Imaging System (BioRad). Only ∼2% of the input SPOR domain protein pelleted in control assay mixtures that lacked PG.

## Acknowledgments

We thank Thomas Bernhardt for the AmiD overproduction plasmid, Simon Foster for the Atl amidase overproduction plasmid, Alex Horswill for strain SH1000 and members of the Weiss lab and Matt Jorgenson for helpful discussions. These studies were supported by NIH GM125656 to DSW, NIH GM138630 to DLP, and NIH R21 AI157121 to JCH. GMK was supported in part by T32AI007511. DNA sequencing was performed at the Genomics Division of the Iowa Institute of Human Genetics which is supported, in part, by the University of Iowa Carver College of Medicine.

